# “*IL1B* loss is associated with increased AR activity in castration-resistant prostate cancer”

**DOI:** 10.1101/2021.08.31.458406

**Authors:** Wisam N. Awadallah, Jagpreet S. Nanda, Sarah E. Kohrt, Magdalena M. Grabowska

**Affiliations:** Department of Urology, Case Western Reserve University, Cleveland, OH; Case Comprehensive Cancer Center, Case Western Reserve University, Cleveland, OH; Department of Pharmacology, Case Western Reserve University, Cleveland, OH; Department of Biochemistry, Case Western Reserve University, Cleveland, OH

**Keywords:** Castration-resistant prostate cancer, androgen receptor, Interleukin 1 beta

## Abstract

Castration-resistant prostate cancer represents a continuum of phenotypes, including tumors with high levels of androgen receptor (AR) expression and activity and those which do not express AR and rely on alternative pathways for survival. The process by which AR-positive prostate cancer cells and tumors lose AR expression and acquire neuroendocrine features is referred to as neuroendocrine differentiation. Numerous therapies and exposures have been demonstrated to induce neuroendocrine differentiation *in vitro*, including the pro-inflammatory cytokine, interleukin 1 beta (IL-1β), encoded by the gene *IL1B*. The purpose of our studies was to determine the relationship between the expression and activity of AR in relationship to IL-1β and *IL1B* in prostate cancer. We performed analysis of de-identified human clinical data and generated prostate cancer cell lines with overexpression or knockout of *IL1B*. In primary prostate cancer, higher expression of *IL1B* predicts longer time to biochemical recurrence. In metastatic castration-resistant prostate cancer, *IL1B* expression is decreased and inversely correlates with *AR* and AR-target gene expression and AR activity, while positively correlating with the neuroendocrine prostate cancer (NEPC) score and neuroendocrine marker gene expression. *In vitro*, we report that AR-positive castration-resistant prostate cancer cells (C4-2B, 22Rv1) secrete IL-1β, and knockout of *IL1B* in these cells results in increased AR activity, in the presence and absence of dihydrotestosterone (DHT). Importantly, knockout of *IL1B* prevented AR attrition during androgen-deprivation. Taken together, our studies demonstrate that loss of *IL1B* in AR-positive castration-resistant prostate cancer cells can increase and maintain AR activity in the absence of androgens, suggesting another potential mechanism of high AR activity in castration-resistant prostate cancer.

## Introduction

Treatment of prostate cancer with androgen deprivation therapy and androgen receptor (AR) signaling inhibitors results in a continuum of castration-resistant prostate cancer phenotypes, including those that maintain high levels of AR, those with neuroendocrine (NE) or small cell features, and those which lack both AR and NE features^1, 2^. The process of epithelial tumor cells acquiring neuroendocrine features has been referred to as “neuroendocrine differentiation”, and reports of this phenomenon in prostate cancer have been reported in pathological studies (reviewed in ^3^) and in response to androgen deprivation therapy/AR inhibition, chemotherapy, and radiotherapy (reviewed in ^4^). While 8-25% of adenocarcinomas can undergo complete neuroendocrine differentiation and become classified as a therapy resistant neuroendocrine prostate cancer ^5–12^, the majority of tumors display focal areas of neuroendocrine differentiation, with up to 85% of castrationresistant prostate cancer patients displaying areas of focal neuroendocrine differentiation^13^.

We have been interested in identifying proteins expressed in these AR-independent areas and tumors, focusing on differential protein expression of transcription factors^14^ and unique adaptations required for telomere maintenance in the absence of AR^15^. In this study, we focused on the pro-inflammatory cytokine interleukin 1 beta (IL-1β), which has been implicated in promoting neuroendocrine differentiation *in vitro*^16, 17^. IL-1β has also been implicated in mediating skeletal colonization^18^ and migration^19^ in AR-independent prostate cancer cell lines, and IL-1β derived from AR-independent prostate cancer cells induces an inflammatory phenotype in bone marrow adipocytes^20^. In skeletal metastatic castration-resistant prostate cancer samples, there is an inverse correlation between prostate specific antigen (PSA) and IL-1β expression^18^.

Interestingly, in primary prostate cancer, higher IL-1β expression in prostate cancer stroma predicts longer time to biochemical recurrence^21^. On a genetic level, there have been several *IL1B* single-nucleotide polymorphisms (SNPs) reported which are associated with increased risk of biochemical recurrence following surgergy^22^, prostate cancer aggressiveness^23^, or risk of developing prostate cancer^24^. Unfortunately, these SNPs have largely not been categorized for impact on IL-1β expression in prostate cancer, and other studies examining genetic polymorphisms of *IL1B* have not found an association with prostate cancer risk^25^. Similarly, other studies have failed to observe relationships between IL-1β expression and biochemical recurrence^26^. *IL1B* expression is higher in prostate cancer samples from African American men versus European American men^27^.

Previous molecular studies have largely focused on the consequences of treatment of prostate cancer cells with exogenous IL-1β^16, 17, 19, 28^, or focused on dissecting IL-1β mediated pathways in AR-independent cell lines^17, 18, 29^. The goal of our study was to rectify these different observations, mainly, is there a functional relationship between AR activity and IL-1β, and can AR-positive castration-resistant prostate cancer cell lines secrete this cytokine, thus supporting AR-independent prostate cancer growth. To do this, we examined publically available clinical data and manipulated *IL1B* in prostate cancer cell lines to examine the consequences on IL-1β secretion and AR signaling.

In clinical samples, *IL1B* expression decreases between primary and metastatic castration-resistant prostate cancer, and is inversely associated with *AR*, AR-target gene expression, and AR activity. Importantly, prostate cancer patients with higher levels of *IL1B* have a longer time to biochemical recurrence. On a molecular level, we report that C4-2B and 22Rv1 cells secrete IL-1β. Knockout of *IL1B* in AR-positive castration resistant prostate cancer cells results in increased AR activity in response to dihydrotestosterone (DHT) treatment. We also report that knockout of *IL1B* results in decreased C4-2B cell line proliferation. Importantly, in the absence of *IL1B*, C4-2B cells are able to maintain higher levels of AR expression under androgen-deprived conditions. Taken together, our results suggest that the loss of IL-1B expression by a subset of castration-resistant prostate cancer patients could support maintained AR expression and activity under androgen-deprived conditions.

## Methods

### Clinical data

For analysis of *IL1B* expression during prostate cancer progression, normalized gene expression data (PCTA Expression data), metadata (PCTA Metadata), and Gleason score data (PCTA PCS category and Gleason score data) were downloaded from the Prostate Cancer Transcriptome Atlas (http://www.thepcta.org/; PCTA Version 1.0.1-Public on February 2, 2021) generated by You et al^30^. These data included 794 benign (noncancer) prostate samples, prostate cancer samples (328 Gleason < 7; 530 Gleason = 7; 203 Gleason > 7), and 260 metastatic castration-resistant prostate cancer samples. Data was integrated using SPSS Statistics 27 (IBM). Gene expression data for *IL1B*, *AR*, AR target genes (*KLK3*, *KLK2*, *FKBP5*), NEPCa markers (*SYP*, *CHGA*, *ENO2*, *FOXA2*), and macrophage markers (*CD68*, *CD86*) was also included. In survival analysis, metastatic castration-resistant prostate cancer cases were excluded. For analysis of the correlation between *IL1B* expression and AR and NEPC activity scores in 208 metastatic castration-resistant prostate cancer samples, data from Abida *et al*^31^ was downloaded from cBioPortal^32, 33^ and reanalyzed following exclusion of one outlier sample.

### Cell culture

Cell lines were cultured as previously described^14^ and parental cell lines underwent short tandem repeat (STR) validation prior to use, along with mycoplasma testing using the LookOut Mycoplasma PCR Detection Kit (Sigma).

### Stable Expression of GFP-tagged IL-1β in LNCaP cells

Lentiviral GFP tagged vector (Origene #100093) and human IL-1β (Origene #RC202079L4) plasmid constructs were used for generating stable LNCaP cells. Packaging plasmid, psPAX2 (Addgene # 12260) and envelope plasmid pMD2.G (Addgene # 12259) were used for lentivirus generation. HEK293FT cells were seeded in 10 cm culture dishes and incubated in DMEM containing 10% FBS overnight (till 70 to 80% confluence). Next day, cells were transfected with either 5μg of GFP tagged empty vector or human IL-1β plasmid along with 3μg of psPAX2 and 2ug of pMD2.G using 30μl of lipofectamine 2000 reagent (Invitrogen, 11668-019). 48 hours post transfection lentivirus enriched medium was collected by centrifugation at 2000 rpm for 10 min and filtered through 0.45 μm sterile syringe filters (VWR International, 28145-481). LNCaP cells in 60mm dishes (0.4 million cells/dish) were infected with 4mL of filtered lentiviral supernatant (diluted 2 times with RPMI1640 medium containing 10% FBS). Two sequential rounds of lentiviral infection were performed in presence of 4.0 μg/mL of hexadimethrine bromide (Sigma, H9268). 48 hours post transfection; cells were re-plated in puromycin containing RPMI 1640 (10% FBS) medium in 10 cm dishes. 1 μg/mL of puromycin was used to select stable LNCaP cells. Whole cell extracts from stable cells were checked for GFP IL-1β expression by western blotting using GFP antibody (ab13970, chick antibody, 1:5000, Abcam).

### Guide RNA design and cloning

Guide RNAs against human IL-1β coding gene were designed using the open access guide RNA design tool (crispr.mit.edu.) developed by the Zhang Laboratory (Broad Institute of MIT and Harvard, McGovern Institute for Brain Research, and Department of Brain and Cognitive Sciences and Biological Engineering, Massachusetts Institute of Technology, Cambridge, MA). Based on higher efficiency score and lower off target events, four guide RNAs (targeting exon 2 or exon 3 of IL-1β) were selected. These guide RNAs were cloned into Bbs1 site of the puromycin-resistant Cas9 expression vector, pSpCas9 (BB)-2A-Puro (PX459) (Addgene # 62988) by using complimentary oligonucleotide pairs. Guide RNA sequences used to cleave IL-1β were as follows: Guide1-GATCATTTCACTGGCGAGCTTC; Guide2-GTGAAATGATGGCTTATTAC; Guide3-GTGATGGCCCTAAACAGATGA and Guide4-GACAGATGAAGGTAAGACTAT. All the recombinant CRISPR Cas9 plasmids with cloned guide sequences were DNA sequenced using U6 forward primer before further use.

### Transient Transfection of C4-2B and 22Rv1 cells for *IL1B* knockout

0.5×10^6^ C4-2B or 0.65×10^6^ 22Rv1 cells seeded overnight in 60mm culture plates were transiently transfected. A combination of CRISPR Cas9 plasmids with guide RNA 2 (targets exon2) and guide RNA 4 (targets exon3) was used for IL-1β knockout in cells. 5ug of empty or guide RNA cloned puromycin-resistant CRISPR Cas9 plasmids (2.5ug of each guide RNA plasmid; wild type control cell line) in Opti-MEM medium with 15 μl of Lipofectamine 2000 was used for each transfection. Medium was changed 8 hours after transfection. Puromycin at a final concentration of 2μg/mL was used for selection for next 2 days (after 48 hours of transfection). Later, cells were trypsinized, counted, seeded in 10 cm dishes (500-1000 cells/dish), and incubated (left undisturbed) for next 2 weeks for colony formation.

### Isolation and screening of *IL1B* knockout clones to generate monoclonal cell lines

Depending upon different growth rates of C4-2B and 22Rv1 cells within 10-16 days of seeding, single isolated colonies were observed under a light microscope. Autoclaved (trypsin soaked) Whatmann filter discs were used to transfer each isolated colony into wells of 48 well tissue culture plates and incubated for 2 weeks. Each confluent well (single clone) was trypsinized and split into two 24 well plates (2 replica plates) which were incubated for next 7-10 days. One of the plates was used for quick genomic DNA isolation for each of the colonies using QuickExtract DNA extraction solution (Lucigen # QE09050). 5μl of this lysate was used to set up a quick genomic PCR screening for IL-1β knockout clones using checking primers. Putative IL-1β knockout clones had an amplicon size ~300bp versus 937bp as compared to the wild type clones or vector controls. Another plate was used to mark these putative clones, expanded and cryopreserved for future use. Putative IL-1β clones were even confirmed by doing western for IL-1β using IL-1β antibody (monoclonal 3ZD anti-IL-1B antibody from NCI). Colonies (wells) which did not show expected IL-1β bands as compared to the vector control on western blotting were confirmed IL-1β knockout clones. Aforementioned single clone cultures were STR validated and screened for any mycoplasma contamination using PCR based Myco detection kit (Sigma, MP0035-1KT) before use. DNA sequences for quick genomic PCR screening primers of IL-1β clones were as follows: Checking primer forward-5’AAAGCCTCTGCTCCAGCTCTCC 3’; Checking primer reverse-5’ AGAGCCCTTCCTTGGGTTGGGA 3’.

### Cell proliferation assays

Cell proliferation in LNCaP *IL1B* overexpression and vector cells was assessed using Cell Proliferation Reagent WST-1 (Roche), per manufacturer’s instructions. Cell proliferation in C4-2B wild type and knockout cells was assessed using the crystal violet assay, as previously described^34^, using 10,000 C4-2B cells/well in a 96-well plate with eight biological replicates per subline. Cell growth was measured on Days 0, 3, 7, and 14.

### Conditioned media analysis by ELISA

To evaluate IL-1B secretion, 10^6 cells were plated in 10 cm tissue culture dish in complete media and then switched to 4mL serum-free, phenol-free RPMI 1640 + L-glutamine (Gibco) for all cell types. After 3 days, cells and conditioned media were collected. Equal amounts of conditioned media were analyzed by ELISA (IL-1 beta Human ELISA Kit, Thermofisher KHC0011), using manufacturer protocol. For data analysis, the average reading from the blank (0.0 pg/mL IL-1β) was subtracted from all values (standards and experimental). A four-parameter sigmoidal standard curve was generated to calculate IL-1B concentrations in conditioned media samples.

### Western blotting

Western blotting was performed as previously described^14^, with the following additions. To evaluate changes in AR under androgen-deprived conditions, 1.0 × 10^6 C4-2B cells were plated in complete media, and the following day were either collected (Day 0) or transferred to serum-free, phenol-free RPMI 1640 + L-glutamine for 7 or 14 days (media was changed at day 7 for the 14 day treatment). Antibodies used include those directed against: IL-1β (clone 3ZD, provided by the BRB Preclinical Repository of the NCI), AR (ab74272, Abcam), and GAPDH (AM4300, Invitrogen).

### Luciferase reporter assays

C4-2B *IL1B* WT, *IL1B* KO clone 3, 5, or 7 cells were plated at 70,000 cells per well of a 24-well plate. 22Rv1 *IL1B* WT, *IL1B* KO clone 8, 10, or 17 cells were plated at 150,000 cells per well of a 24-well plate. After 24 hours, cells were transfected with 0.5μg/well of the composite probasin promoter ARR_2_PB-luciferase^35, 36^ or prostate-specific antigen enhancer-promoter construct PSAEP-luciferase^37^ and 0.05μg/well SV40-renilla constructs (transfection control) in Lipofectamine 2000 in charcoal-stripped media (phenol-free RPMI 1640 with L-Glutamine plus 10% charcoal-stripped FBS) for 24 hours. Cells were then induced with 10nM ethanol (vehicle control) or dihydrotestosterone for 24 hours. Luciferase and renilla expression was quantified using the Dual-Reporter assay system (Promega) using a SpectraMax ID3 plate reader with a dual injection system (Molecular Devices). Luciferase data was then normalized to renilla and then vector cells.

### Statistical analysis

Data was analyzed in SPSS and GraphPad Prism and analyzed using parametric or non-parametric tests depending on normality of the data distribution.

## Results

To determine the expression of *IL1B* during prostate cancer progression, we leveraged the clinically annotated and normalized gene expression data from the Prostate Cancer Transcriptome Atlas (http://www.thepcta.org/) generated by You et al^30^. These data included 794 benign (non-cancer) prostate samples, 1,361 prostate cancer samples (328 Gleason < 7; 530 Gleason = 7; 203 Gleason > 7), and 260 metastatic castration-resistant prostate cancer samples. We first compared *IL1B* expression between benign, prostate cancer, and metastatic castration-resistant prostate cancer. While there was no statistically significant difference in *IL1B* expression between primary prostate cancer and benign prostate tissue, *IL1B* expression was decreased in metastatic castration-resistant prostate cancer samples versus primary (P < 0.001, Figure 1A, next page). As IL-1β has been reported in bone metastases from prostate cancer patients, we also compared *IL1B* expression between different metastatic locations of castration-resistant prostate cancer patients and report that metastatic samples from the bone had higher expression of *IL1B* versus castration-resistant prostate cancer samples from the prostate (P < 0.01, Figure 1B).

**Figure 1:**
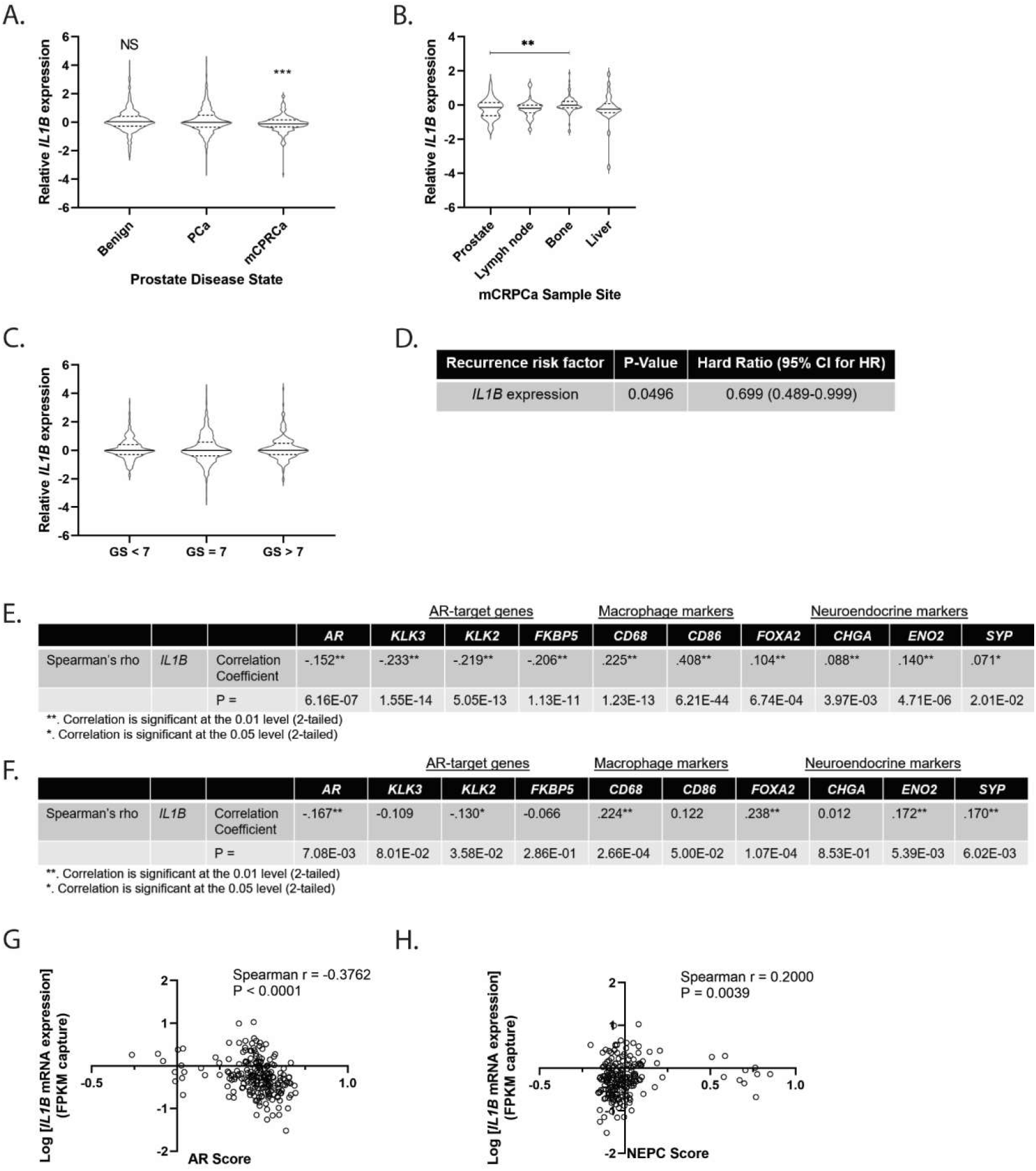
*IL1B* expression decreases during progression to castration-resistance and correlates with a neuroendocrine prostate cancer gene signature. **A.** *IL1B* expression decreases in metastatic castration-resistant prostate cancer. Analysis of normalized *IL1B* expression in benign (non-tumor, n = 794), prostate cancer (PCa, n = 1061), and metastatic castration-resistant prostate cancer (mCRPCa, n = 260). Comparisons made versus prostate cancer, with Kruskal-Wallis test with Dunn’s multiple comparisons. **B.** *IL1B* expression is increased in bone metastases versus prostate-derived metastatic castration-resistant prostate cancer samples. Analysis of normalized *IL1B* expression in castration-resistant prostate cancer samples derived from prostate (n = 54), lymph node (n = 57), bone (n = 86), and liver (n = 30). Comparisons made versus prostate, with Kruskal-Wallis test with Dunn’s multiple comparisons. **C.** *IL1B* expression is not affected by Gleason grade. Analysis of Gleason Score (GS) tumors less than 7 (GS < 7, n = 328), equal to 7 (GS = 7, n = 530), and greater than 7 (GS > 7, n = 203). Comparisons made versus GS < 7, with Kruskal-Wallis test with Dunn’s multiple comparisons. **D.** Increasing *IL1B* predicts longer time to recurrence. Cox regression analysis of *IL1B* association with time to recurrence in 131 patients (77 with recurrence, 54 censored/without). HR: Hazard Ratio; CI: Confidence Interval. **E/F**. *IL1B* expression positively correlates with neuroendocrine differentiation genes and negatively correlates with AR and AR-target gene expression in primary (E) and metastatic castration-resistant prostate cancer (F). Spearman correlation analysis between *IL1B* and AR, AR target genes (*KLK3*, *KLK2*, *FKBP5*), neuroendocrine differentiation/neuronal genes (*FOXA2*, *CHGA*, *ENO2*, *SYP*) and macrophage genes (*CD68*, *CD86*) in 1061 prostate cancer (E) or 260 metastatic castration-resistant prostate cancer samples (F). **G.** *IL1B* expression inversely correlates with AR activity in metastatic castration-resistant prostate cancer. Spearman correlation analysis between logIL1B and AR activity (n = 207). **H.** *IL1B* expression correlates with a neuroendocrine prostate (NEPC) gene signature in metastatic castration-resistant prostate cancer. Spearman correlation analysis between logIL1B and AR activity (n = 207) NS: Not Significant; ** P < 0.01; *** P < 0.001

Previously, stromal IL-1β expression was reported to be protective against prostate recurrence; meanwhile tumor-specific IL-1β expression did not achieve statistical significance for this effect (Hazard Ratio [HR]: 0.431, [95% Confidence Interval (CI): 0.180–1.029], P = 0.058)^21^. To evaluate whether increased *IL1B* expression was predictive of longer time to biochemical recurrence, we first evaluated whether *IL1B* expression was similar across disease grade. Consistent with observations that IL-1β expression is uniform between Gleason grades^21^, we did not observe changes in *IL1B* expression between Gleason grades (Figure 1C). We then performed Cox regression analysis on the 131 patients with recurrence data (77 with recurrence, 54 censored/without) and found that increasing levels of *IL1B* expression was associated with longer time to biochemical recurrence (HR: 0.699 [95% CI: 0.489-0.999], P = 0.0496, Figure 1D).

As IL-1β has been reported to induce neuroendocrine differentiation of prostate cancer cell lines *in vitro*^16, 17^, we next analyzed the correlation between *IL1B* expression and genes associated with neuroendocrine differentiation (*FOXA2*, *CHGA*, *ENO2*, *SYP*), AR, and AR-target genes (*KLK3*, *KLK2*, *FKBP5*) in primary and metastatic castration resistant prostate cancer. As IL-1β is largely expressed by macrophages and tumor samples contain a mixture of different cell types (tumor, stroma, immune) we also examined macrophage markers (*CD68*, *CD68*) to see whether changes in *IL1B* were associated with a proxy for macrophage presence. In primary prostate cancer, *IL1B* was negatively correlated with *AR*, *KLK3*, *KLK2*, and *FKBP5* expression (Figure 2E). In these samples, *IL1B* was positively correlated with *FOXA2*, *CHGA*, *ENO2*, *SYP*, *CD86*, and *CD68*. In metastatic castration-resistant prostate cancer, *IL1B* was negatively correlated with *AR* and *KLK2*, and positive correlated with *FOXA2*, *ENO2*, *SYP*, and *CD68* (Figure 1F). To move beyond a gene-to-gene correlative analysis, we also analyzed AR activity and similarity to a neuroendocrine prostate cancer (NEPC) profile in metastatic castration-resistant prostate cancer samples from Abida *et al^31^*. Indeed, *IL1B* expression inversely correlates with AR activity score (Figure 1G) and positively correlates with NEPC score (Figure 1H).

**Figure 2:**
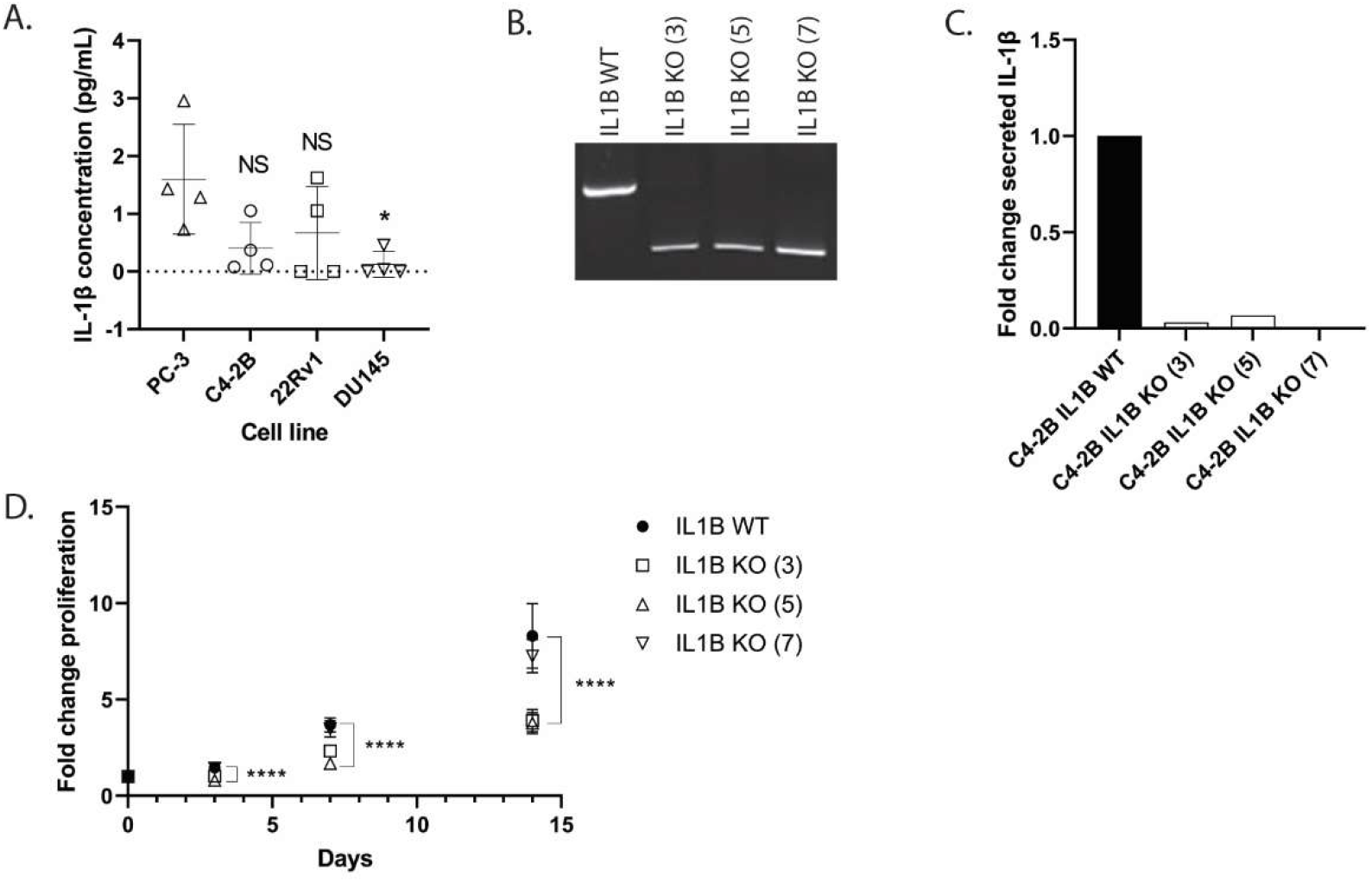
Knockout of *IL1B* decreases proliferation in C4-2B cells. **A.** IL-1B is secreted by PC-3, C4-2B, and 22Rv1 cells. ELISA analysis of IL-1β secretion into media by prostate cancer cell lines. Statistical comparison to PC-3 cells (positive control) via Kruskal-Wallis test with Dunn’s multiple comparisons. **B.** *IL1B* knockout in C4-2B cells. PCR confirmation of *IL1B* knockout spanning the CRISPR/Cas9 excision site in *IL1B* knockout (*IL1B* KO (3), *IL1B* KO (5), *IL1B* KO (5), ~300bp amplicon) monoclonal cell lines with the *IL1B* wild type (*IL1B* WT, 937bp amplicon) monoclonal cell line serving as a control. **C.** *IL1B* knockout cells have a dramatic reduction in IL-1B secretion. Fold change secreted IL-1β as measured by ELISA analysis of wild type and *IL1B* knockout (monoclonal cell line 3, 5, and 7) conditioned media. **D**. Knockout of *IL1B* can decrease C4-2B proliferation. Fourteen day cell proliferation analysis of C4-2B sublines, with analysis by ordinary one-way ANOVA with Dunnett’s multiple comparisons test. Significant statistical differences denoted between wild type (WT) and knockout (KO) cell lines clone 3 and 5. NS: Not Significant; **** P < 0.0001

Based on these observations, we next set out to evaluate whether endogenous IL-1β produced by cancer cells impacts prostate cancer cell growth. We therefore generated *IL1B* overexpressing and knockout cell lines in androgen-dependent and castration-resistant prostate cancer cell lines, respectively. Consistent with our observation in clinical data, when we overexpressed *IL1B* in androgen-dependent LNCaP cells, we observed a decrease in growth (Supplemental Figure 1A, B, final page) to the point that we were unable to maintain stable *IL1B* overexpressing cells long-term. This is consistent with previous reports that demonstrated LNCaP cells do not express *IL1B*^38^ and where exogenous IL-1β treatment of LNCaP cells resulted in decreased proliferative capacity^16^. Due to this limitation, and that our analysis indicated that *IL1B* expression was decreased in metastatic castration-resistant prostate cancer, we focused on castration-resistant prostate cancer cell lines.

Previous reports demonstrated that PC-3 cells secrete IL-1B^18^, while DU145 have low^18^ or undetectable^38^ IL-1β levels. In serum free conditioned media, we observed PC-3 cells secreting 1.6 pg/ml, consistent with other reports of ~2 pg/ml^18^. There was no statistically significant difference between PC-3, 22Rv1, and C4-2B IL-1β secretion (Figure 2A), but DU145 cells did secrete significantly less (0.1235 pg/ml). This observation was supported by western blotting from whole cell lysates that demonstrated mature (cleaved) IL-1β in PC-3, C4-2B, and 22Rv1 cells, with low levels in DU145 cells (Supplemental Figure 1C).

To determine whether the loss of endogenous IL-1β impacted AR-positive castration-resistant prostate cancer cell lines, we generated monoclonal *IL1B* knockout C4-2B and 22Rv1 cells. We confirmed *IL1B* knockout using PCR spanning the CRISPR/Cas9 targeted sequence and identified three knockout monoclonal cell lines (sublines) in each cell line for use (Figure 2B, Supplemental Figure 2A). We also confirmed *IL1B* knockout by measuring changes in IL-1β secretion in C4-2B cells using ELISA (Figure 2C). We then examined the growth of *IL1B* wild type and knockout C4-2B sublines over a 14-day time course. Surprisingly, two of three knockout sublines had decreased proliferation versus wild type control cells (Figure 2D).

In our analysis of clinical data, we observed an inverse correlation between AR activity and *IL1B* expression. To determine whether this correlation could be driven by tumor cells, rather than tumor-associated macrophages, we examined AR activity in *IL1B* wild type C4-2B (C4-2B-*IL1B-WT*) and *IL1B* knockout (C4-2B-IL1B-KO) cells. Indeed, knockout of *IL1B* in C4-2B cells resulted in a dramatic increase in AR activity in response to DHT from the ARR_2_PB-luciferase^35^ reporter construct in C4-2B-IL1B-KO (3) and C4-2B-IL1B-KO (7) cell lines versus control cells (Figure 3A, next page). In the C4-2B-IL1B-KO (5) cell line, increased AR activity was observed in some experiments, but not others. When we analyzed AR activity using the PSAEP-luciferase^37^ construct, only the C4-2B-IL1B-KO (7) cell line exhibited increased AR activity in response to DHT (Figure 3B). Importantly, in the absence of androgens, knockout cells had higher AR activity at baseline (Figure 3A, B). In 22Rv1 cells, we observed a similar pattern of increased AR activity (Supplemental Figure 2B, C).

**Figure 3:**
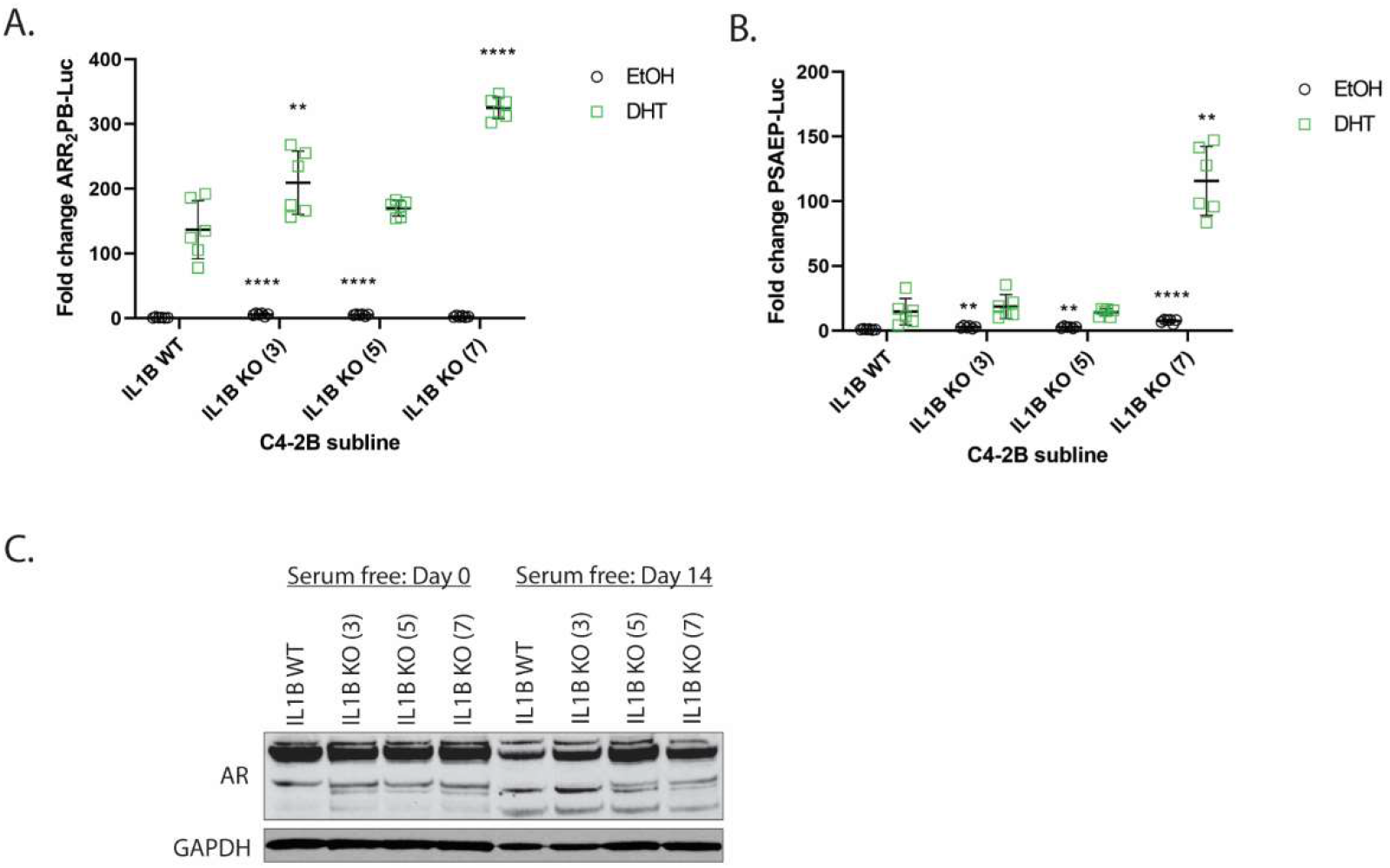
Knockout of *IL1B* increases AR activity in C4-2B cells. **A/B**. Knockout of *IL1B* increases AR activity. Comparison of wild type and *IL1B* knockout cells for changes in AR activity in response to dihydrotestosterone (DHT) or vehicle (ethanol, EtOH), as measured by ARR_2_PB (D) and PSAEP (E) luciferase activity. Data normalized to renilla and then wild type vector control, with analysis by ordinary one-way ANOVA with Dunnett’s multiple comparisons test. Statistical differences denoted between vehicle-to-vehicle and DHT-to-DHT comparisons. **C.** Knockout of *IL1B* results in maintained AR expression under androgen-deprived conditions. Comparison of wild type and knockout cell AR expression after 0 or 14 days of growth in serum free media. WT: Wild Type; KO: Knockout; NS: Not Significant; * P < 0.05; ** P < 0.01; *** P < 0.001; **** P < 0.0001

As IL-1β has been associated with neuroendocrine differentiation, *IL1B* expression is associated with neuroendocrine prostate cancer gene expression (Figure 1), and exogenous IL-1β treatment of C4-2B cells can induce neuroendocrine differentiation^17^, we decided to focus on C4-2B cells and compare wild type control cells to knockout lines C4-2B-IL1B-KO (3) and C4-2B-IL1B-KO (7). In previous reports IL-1β treatment of C4-2 cells, a precursor of C4-2B cells, reduced *AR* expression^28^, but whether loss of endogenous IL-1β could prevent AR loss had not been evaluated. We therefore challenged control and knockout cells with serum free media for 0 and 14 days to examine whether loss of *IL1B* prevented loss of AR. At baseline, C4-2B-IL1B-WT cells express equal amounts of AR versus C4-2B-IL1B-KO cells. Following 14 days of serum-free media treatment, we observed that C4-2B-IL1B-WT cells had lower AR expression than C4-2B-IL1B-KO cells (Figure 3C).

## Discussion

The goal of these studies was to evaluate whether AR-positive castration-resistant prostate cancer cell lines secrete IL-1β and determine the consequences of inhibiting IL-1β secretion on AR activity. We report that castration-resistant prostate cancer cells (C4-2B and 22Rv1) secrete IL-1β at levels comparable to the AR-independent PC-3 cell line and that loss of *IL1B* reduces the proliferative capacity of C4-2B cells. Consistent with previous studies implicating IL-1β treatment with neuroendocrine differentiation^16, 17^, knockout of *IL1B* maintained higher AR expression in C4-2B cells under serum-deprived conditions. Knockout of *IL1B* also mediated increased AR activity as assessed by AR-reporter assays. These *in vitro* observations are supported by clinical data where there is an inverse relationship between AR activity and *IL1B*.

Our study has several limitations. First, while we are confident that prostate cancer cell lines are capable of secreting IL-1β, our studies have not evaluated the mechanisms by which IL-1β secretion is activated nor how IL-1β is processed to its active form. Furthermore, our studies have not demonstrated which pathways connect changes in IL-1β secretion and AR activity. Our studies also exclusively focus on IL-1β secretion by cancer cells. Mechanistic studies addressing these unanswered questions represent important next steps to understanding the interplay between immune cells, cytokine secretion, and castration-resistant prostate cancer cells.

Another limitation of our study is that we do observe some variability in our monoclonal knockout cell lines in terms of proliferation and AR activity response. This likely has to do the with variability in monoclonal cell lines generated by CRISPR/Cas9 arising from transfection of different cell line subpopulations, off-target effects of CRISPR/Cas9, and the monoclonal cell line selection process. This variation was observed in two castrationresistant prostate cancer cell lines (C4-2B, 22Rv1) and suggests that the response to *IL1B* loss could be contextdependent in patient tumors. This observation fits into our clinical data analysis, where the correlations between *IL1B* expression and AR activity and the NEPC score are statistically significant but not associated with large effect sizes.

Our studies fit into a general model where tumors with focal neuroendocrine differentiation have two cancer cell types: those with and without AR. We propose that IL-1β secreting AR-positive castration-resistant prostate cancer cells can support AR-negative cell line growth and neuroendocrine differentiation, thus leading to a neuroendocrine or AR-negative phenotype. Conversely, if IL-1β secretion in AR-positive castration-resistant prostate cancer cells is prevented, this results in increased AR activity and could result in increased outgrowth of castration-resistant prostate cancer cells with high AR activity, even in the absence of androgens.

## Acknowledgements

This work was supported by R00 CA197315 (to MMG). The content of this manuscript is solely the responsibility of the authors and does not necessarily represent the official views of the NIH. We would also like to acknowledge the Case Research Institute, a joint venture between University Hospitals and Case Western Reserve University, start-up funds (to MMG), and the Molecular Therapeutics Training Program (MTTP) (T32 GM 008803 to SEK).

---

**Supplemental Figure 1:**
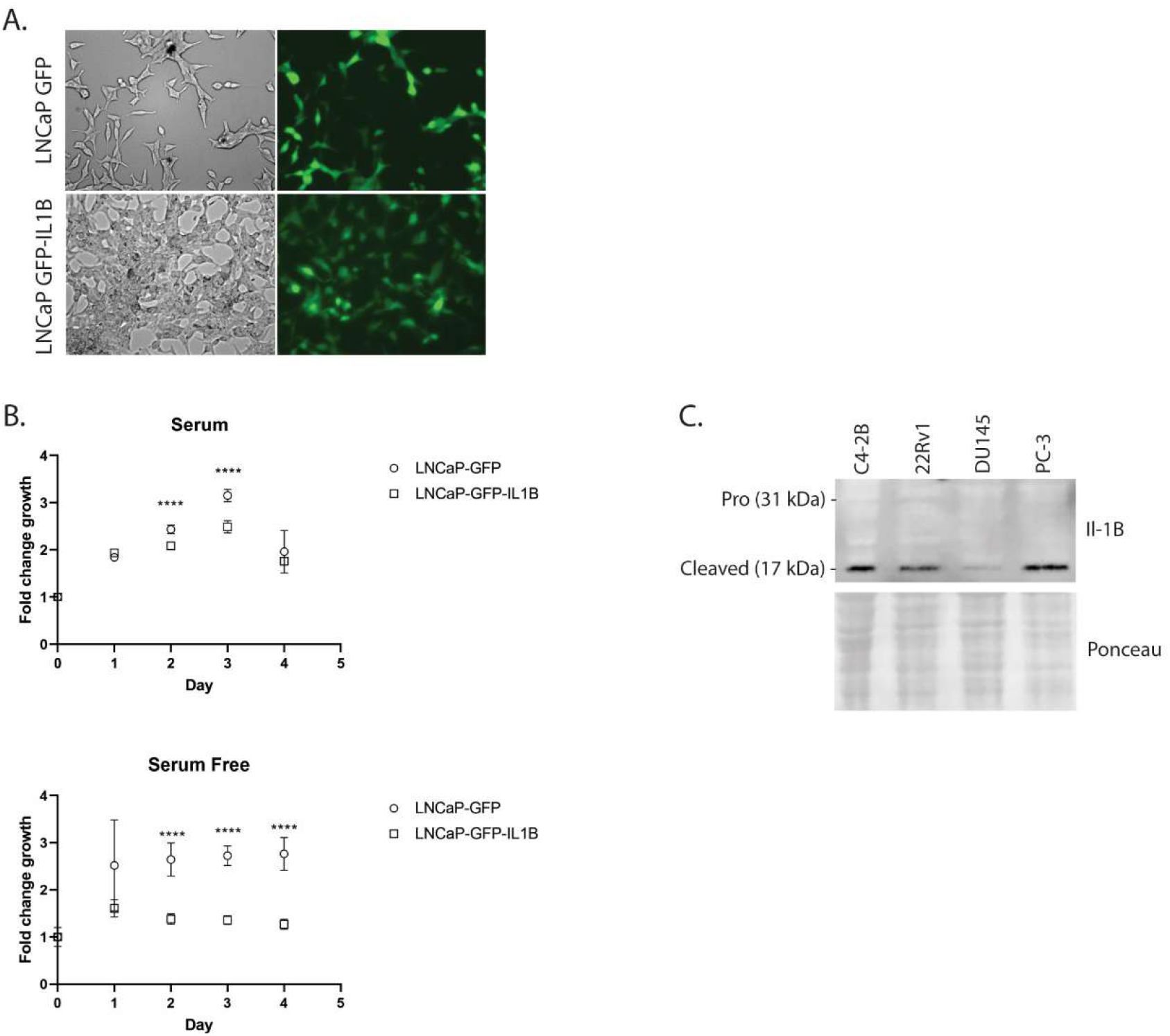
IL-1β expression in prostate cancer cells. **A.** Generation of *IL1B* overexpressing LNCaP cells. GFP and GFP-IL1B overexpressing LNCaP cells. **B.** *IL1B* overexpression in LNCaP cells reduces cell proliferation. Proliferation of LNCaP-GFP and LNCaP-GFP-IL1B in serum containing and serum free media, measured over four days, normalized to day 0 readings relative to each cell line. Comparisons by unpaired t-test between control and experimental cells at each time point. **** P < 0.0001 **C.** Prostate cancer cell lines express cleaved (mature) IL-1B. Western blot analysis of prostate cancer cell whole cell lysate for IL-1β.

**Supplemental Figure 2:**
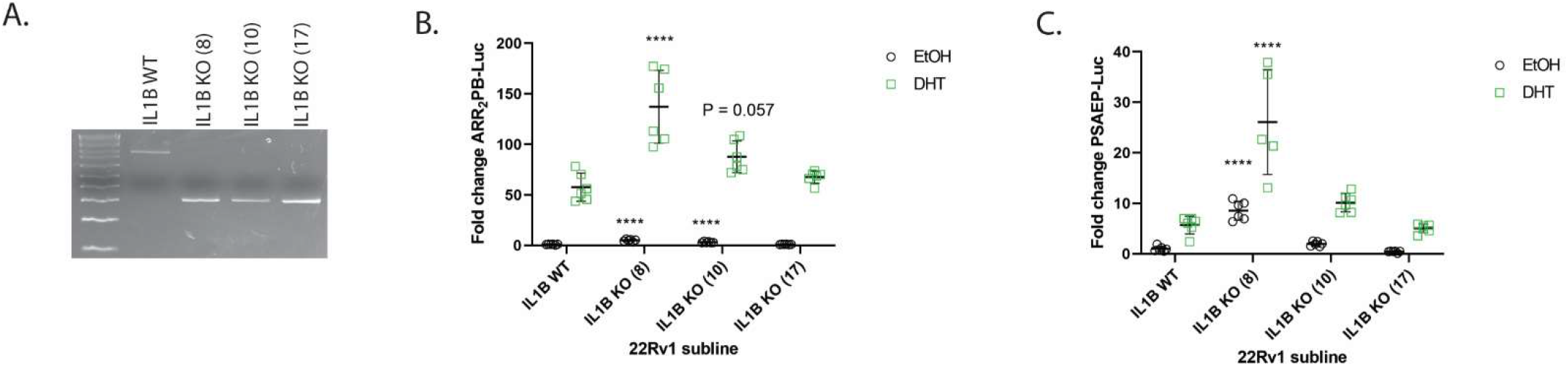
Knockout of *IL1B* increases AR activity in 22Rv1 cells. **A.** *IL1B* knockout in 22Rv1 cells. PCR confirmation of *IL1B* knockout spanning the CRISPR/Cas9 excision site in *IL1B* knockout (*IL1B* KO (8), *IL1B* KO (10), *IL1B* KO (17)) monoclonal cell lines with the *IL1B* wild type (*IL1B* WT) monoclonal cell lines serving as a control. **B/C**. Knockout of *IL1B* increases AR activity in 22Rv1 cells. Comparison of wild type and *IL1B* knockout cells for changes in AR activity in response to dihydrotestosterone (DHT) or vehicle (ethanol, EtOH), as measured by ARR_2_PB (B) and PSAEP (C) luciferase activity. Data normalized to renilla and then wild type vector control, with analysis by ordinary one-way ANOVA with Dunnett’s multiple comparisons test. Statistical differences denoted between vehicle-to-vehicle and DHT-to-DHT comparisons. WT: Wild Type; KO: Knockout; **** P < 0.0001

